# Co-evolution of host dispersal and parasite virulence in complex landscapes

**DOI:** 10.1101/2024.10.28.620220

**Authors:** Jhelam N. Deshpande, Ruthvik S. Pallagatti, Vasilis Dakos, Oliver Kaltz, Emanuel A. Fronhofer

## Abstract

Spatial network structure impacts the ecological and evolutionary dynamics of species interactions. Previous work on host-parasite systems has shown that parasite virulence is driven by dispersal rates and spatial structure, assuming that dispersal is an ecologically fixed parameter. However, dispersal is also a trait under selection and can evolve. In this context, we develop an individual-based eco-evolutionary model, in which both parasite virulence and host dispersal can evolve in representative terrestrial (random-geometric graphs; RGGs) and riverine aquatic (optimal channel networks; OCNs) landscapes. We find that in riverine aquatic landscapes, evolutionarily stable (ES) dispersal rates are lower and ES virulence is greater relative to terrestrial landscapes when dispersal mortality is low. When dispersal mortality is high, both dispersal and virulence evolve to lower values in both landscape types. Diverging co-evolutionary patterns between landscapes are explained by differences in network topology. Specifically, the highly heterogeneous degree distribution in riverine aquatic landscapes 1) leads to low parasite relatedness allowing for the evolution of greater virulence and 2) leads to spatial heterogeneity in host densities that constrains the evolution of dispersal to lower values. Our work highlights the importance of considering the concurrent and co-evolution of dispersal when studying trait evolution in complex landscapes.

## Introduction

Ecological systems consist of networks of interactions, such as food webs, host-parasite networks, or plant-pollinator networks, that are embedded in spatial networks, thereby forming meta-networks (Melián et al., 2018). Importantly, landscapes may be shaped in various configurations, in water or on land, producing specific types of networks with variable topology (Urban and Keitt, 2001). These idiosyncrasies of spatial network structure can impact both ecological and evolutionary dynamics, as well as eco-evolutionary feedbacks (Govaert et al., 2019; Fronhofer et al., 2023). This may have profound effects on the spatial extent of biotic interactions and their evolutionary dynamics. Thus, if one wants to understand the evolution of traits governing interactions between species, such as competitive ability, predation rate, or virulence, we need to explicitly account for the landscape context.

Further, the impact of spatial network structure on ecological and evolutionary dynamics of interacting species cannot be understood without accounting for the key role dispersal rates and distances play within landscapes by defining local demography and gene flow (Govaert et al., 2019; Fronhofer et al., 2023). Moreover, dispersal is a trait that has a genetic basis (Saastamoinen et al., 2018) and may itself evolve (Ronce, 2007) in response to spatial network structure (Henriques-Silva et al., 2015; Fronhofer and Altermatt, 2017). As a consequence, to gain a complete picture of how traits mediating interactions between species evolve in meta-networks, one needs to consider the co- and concurrent evolution of traits defining species interactions with dispersal (Zilio et al., 2024).

Here, we focus on the impact of spatial network structure on the co-evolution of parasite virulence and host dispersal. Host-parasite systems are ubiquitous, and it is therefore important to understand important parasite traits, such as virulence (i.e., the extent to which a parasite harms its host). Since host-parasite systems are often spatially structured (Parratt et al., 2016; Penczykowski et al., 2016; White et al., 2018), spatial aspects of virulence evolution have been studied extensively in theoretical models (Messinger and Ostling, 2009; Lion and Gandon, 2015). In general, a number of models (Boots and Sasaki, 1999; O’Keefe and Antonovics, 2002; Wild et al., 2009; Lion and Boots, 2010) have highlighted the central role of kin selection: In spatially structured systems, lower parasite virulence evolves as an altruistic strategy such that parasites leave a larger pool of susceptible hosts to related individuals. Yet, most studies assume simplified spatial structures, such as lattice or island models. In complex spatial networks, such as those modelling terrestrial (Gilarranz, 2020) or riverine aquatic landscapes (Rinaldo et al., 2014; Carraro et al., 2020), parasite virulence evolution is also driven by kin selection, with the two landscape types producing characteristic patterns of parasite relatedness and host dispersal that in turn produce distinct relationships between parasite virulence and host dispersal (Deshpande et al., 2023). More specifically, virulence is predicted to be unimodal as a function of host dispersal, attaining a higher maximum in riverine aquatic landscapes, and is a saturating function of host dispersal in terrestrial landscapes (Deshpande et al., 2023).

Thus, the precise effect of landscape structure on evolved parasite virulence depends on assumed host dispersal rates (Deshpande et al., 2023). However, as mentioned above, dispersal can also evolve in response to landscape structure (Henriques-Silva et al., 2015; Fronhofer and Altermatt, 2017). Indeed, the mechanisms driving the evolution of dispersal in a single species context have been studied extensively (Duputié and Massol, 2013; Bonte et al., 2012; Bowler and Benton, 2005). Generally, increased dispersal evolves in response to spatio-temporal variation in fitness expectation, which can result from kin competition, demographic stochasticity, and extinction-recolonisation dynamics. Conversely, dispersal evolution may be constrained by associated costs such as reduced survivorship during travel (Bonte et al., 2012) or if there is temporally invariant spatial heterogeneity in fitness expectation (Hastings, 1983). These basic mechanisms also explain dispersal evolution in complex spatial networks. Fronhofer and Altermatt (2017) show that larger average landscape connectivity increases dispersal through its effect on kin structure, hence kin competition, while larger heterogeneity in connectivity, typical of riverine landscapes or scale-free networks, generates temporally invariant heterogeneity in population density, which selects against dispersal (Henriques-Silva et al., 2015).

Therefore, in order to fully understand eco-co-evolution in meta-networks, one must study the evolution of both dispersal and interaction traits simultaneously. This is especially true since we can expect feedback between such co-evolving traits: As Zilio et al. (2024) discuss, the evolution of interaction traits in antagonistic species strongly impacts population densities and the spatial distribution of interaction partners. This, in turn, directly impacts the various mechanisms described above that drive the evolution of dispersal, which remain unchanged from a single species context (Fronhofer et al., 2023). Previous work focusing on dispersal evolution alone in host-parasite (Green, 2009; Chaianunporn and Hovestadt, 2012b) and predator-prey systems (Pillai et al., 2012; Amarasekare, 2016) has emphasised the key role of extinction-recolonisation dynamics that depend on the strength of interactions between antagonistic partners. However, these studies do not consider the co-evolution of interaction traits and assume simplified landscape structures.

Very few studies consider dispersal evolution together with trait evolution. Kamo et al. (2007) study the concurrent evolution of parasite virulence and proportion of global infection (parasite dispersal) and find that parasite fitness is maximised at intermediate dispersal and higher transmission than is expected from well mixed systems. Chaianunporn and Hovestadt (2012a) show that heterogeneity in the abiotic environment of the host selects against dispersal during the concurrent evolution of host/parasite dispersal and niche width. Furthermore, consumer-resource dynamics may produce characteristic patterns of local adaptation that impact dispersal evolution (Drown et al., 2013). Finally, the concurrent evolution of dispersal can promote the transition from parasitism to mutualism (Ledru et al., 2022).

While all of these studies contribute to our understanding of host-parasite eco-co-evolution, none takes the step to study eco-co-evolution in complex landscapes (Zilio et al., 2024). Here, we extend the modelling framework used by Deshpande et al. (2023) to study the co-evolution of host dispersal and parasite virulence in complex landscapes. We focus on two spatial network structures representative of two major biomes worldwide: terrestrial landscapes, which can be represented using random-geometric graphs (RGGs; Gilarranz, 2020) and riverine aquatic landscapes, represented by optimal-channel networks (OCNs; Rinaldo et al., 2014; Carraro et al., 2020). We ask the following question: How does the evolution of host dispersal, mediated by dispersal costs (Bonte et al., 2012), impact the evolution of parasite virulence, and vice versa, in terrestrial vs. riverine aquatic landscapes?

## Model description

### Model overview

In our individual-based model, both host and parasite are asexual, and only host individuals are explicitly represented as either susceptible (do not bear a parasite) or infected (bear a parasite). The model assumes discrete-time with non-overlapping host and parasite generations, and spatial structure is also discrete since we model the metapopulation as a network of *n* = 100 patches. We explore two landscape types: terrestrial, modelled by random-geometric graphs (RGGs; Gilarranz, 2020), and riverine aquatic, modelled by optimal channel networks (OCNs; Rinaldo et al., 2014; Carraro et al., 2020). Hosts disperse after they are born, and individuals may die during dispersal with a probability *µ* representing dispersal costs (Bonte et al., 2012). After dispersal, hosts give birth to susceptible offspring (no vertical transmission) in their target patches and die, after which parasite propagules from the infected hosts of the parental generation may infect individuals of the offspring generation. We model parasite virulence acting on the fecundity of the host and host dispersal probability as quantitative traits that can evolve. We run the model for 5000 time steps until both parasite virulence and host dispersal reach (quasi)-equilibrium to obtain ES trait values. Below we explain in detail the assumed landscape structures, the life cycle of the modelled host and parasite, and the analyses performed.

### Landscape structure

Landscape structure is represented by an adjacency matrix **D**_*n×n*_. Two patches *x* and *y* in the meta-network are connected to each other by dispersal if (*D*_*x*,*y*_) = 1 and they are not connected to each other if (*D*_*x*,*y*_) = 0. Specifically, we model two types of networks:

#### Terrestrial landscapes: Random-geometric graphs (RGGs)

Following Gilarranz (2020), Saade et al. (2023) and Deshpande et al. (2023), we represent terrestrial landscapes using RGGs. The physical process assumed here is as follows: Habitat patches are distributed uniformly on a two-dimensional piece of land (assumed for simplicity to be a square). Coordinates of *n* = 100 patches are drawn from a uniform distribution on the unit square *U* [0, 1] *×* [0, 1], and due to intrinsic biological limitations on dispersal distances, dispersal is possible between two patches only if they are within a radius *r* of each other. Here, we fix *r* = 0.15, which implies an average degree (average number of neighbours a patch has) of approximately 6 (Deshpande et al., 2023).

As opposed to classically used networks, such as grids (Chaianunporn and Hovestadt, 2012b), that are used to represent terrestrial host-parasite systems, these structures are more realistic since they have variation in their connectivity rather than a fixed degree and are modular, which is a characteristic of terrestrial systems (Gilarranz, 2020). We generate an ensemble of 1000 RGGs.

#### Riverine aquatic landscapes: Optimal channel networks (OCNs)

OCNs take into account the geomorphological processes that generate rivers globally (Rinaldo et al., 2014). The OCN concept is based on the assumption that river networks correspond to a minimum of total energy dissipation across the landscape and the generated structures reproduce all scaling features of real river networks throughout the globe. We model an unaggregated river network with a fixed outlet position with 10 *×* 10 patches, and generate their connectivity using the R package OCNet (version 0.5.0; Carraro et al. 2020). These networks have an average degree of 1.98 and are characterised by a highly heterogeneous degree distribution, with many degree-1 patches (i.e., patches with only 1 neighbour) and very few patches with a greater degree. We generate an ensemble of 1000 OCNs.

#### Hexagonal grids and circular landscapes

In order to compare our results to simplified spatial structures, we run additional simulations on regular hexagonal grids (average degree 6, which is close to the average degree of the terrestrial landscapes assumed) and in circular networks (average degree 2, which is close to the average degree of the aquatic landscapes).

### Life cycle

#### Host Dispersal

Host dispersal is natal, i.e., hosts disperse before reproduction, to any neighbouring patch defined by the connectivity matrix with equal probability. Individuals may die during dispersal with a probability *µ* which captures costs associated with dispersal (Bonte et al., 2012).

#### Host reproduction and inheritance

Hosts reproduce after dispersal. Population dynamics of the host are logistic with local density regulation according to the Beverton-Holt model (Beverton and Holt, 1957). Thus, if there are *S*_*x*_(*t*) susceptible individuals, *I*_*x*_(*t*) infected individuals, and a total of *N*_*x*_(*t*) = *S*_*x*_(*t*) + *I*_*x*_(*t*) in a patch *x* post dispersal, the expected population density at time *t* + 1, is given by:

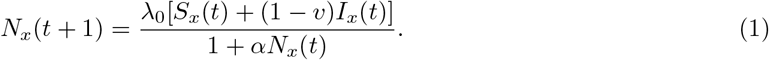

Here, *λ*_0_ is the intrinsic growth rate, *α* is the intraspecific competition coefficient, and 0 ≤ *v* ≤ 1 is the expected parasite virulence acting on the fecundity of the host (Abbate et al., 2015; Chaianunporn and Hovestadt, 2012b; Deshpande et al., 2023). Thus, the expected number of offspring an individual *k* in a patch *x* produces is 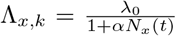 and 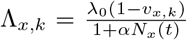 if infected, where *v*_*x*,*k*_ is the genotypic value of virulence of the parasite infecting that individual. Finally, population dynamics are stochastic; therefore, the realised number of offspring for an individual are drawn from a Poisson distribution with mean Λ_*x*,*k*_. All offspring are born susceptible; there is no vertical transmission of the parasite.

These offspring inherit their dispersal trait from their parent. During inheritance, the dispersal trait mutates with a certain fixed probability *m*_*d*_ with mutation effects drawn from a normal distribution with mean 0 and standard deviation and logit transformed standard deviation *σ*_*d*_. After reproduction, the parental host generation dies.

#### Parasite transmission

Once the hosts of the parental generation die, the parasites from dead infected individuals release spores, which may infect individuals from the offspring generation. Thus, the expected dynamics of the number of infected individuals *I*_*x*_(*t* + 1) in a patch *x* at time *t* + 1, which had *I*_*x*_(*t*) parental infected individuals, follow a modified Nicholson-Bailey model (Nicholson and Bailey, 1935) with a type-II functional response (Chaianunporn and Hovestadt, 2012b; Deshpande et al., 2021, 2023), and are given by:

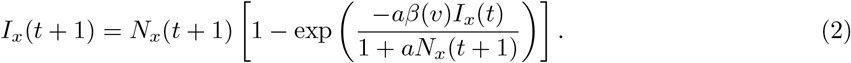

Here, *β*(*v*) is the transmission rate, which is assumed to be an increasing function of virulence, and *a* is the searching efficiency of the parasite. We assume that the more a parasite exploits its host, therefore harming it, the more it can transmit (Alizon et al., 2009), thus,

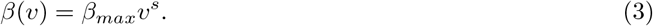

Here, *β*_*max*_ *>* 0 is the maximum transmission rate, which is obtained at *v* = 1, and *s >* 0 determines the shape of this relationship (*s <* 1, *s* = 1, and *s >* 1 implies decelerating, linear, and accelerating relationships between virulence and transmission, respectively).

Therefore, at an individual level, we assume that a propagule released by an infected parent *k* in a patch *x* comes in contact with a newborn susceptible with a probability 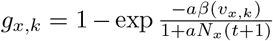. Thus, multiple parasite propagules can come in contact with a given offspring host, but ultimately only one infects it. During transmission, the virulence trait of the parasite is inherited, and it mutates with a fixed probability *m*_*v*_ with mutation effects drawn from a normal distribution centred around 0 with a logit transformed standard deviation *σ*_*v*_ of 0.5 to ensure that the trait remains constrained between 0 and 1.

### Simulations

We ran simulations on 1000 realisations of each landscape type for 5000 generations, which allowed the populations to reach a quasi-equilibrium (Fig. S1) and obtain ES trait values. The simulations were initiated with 300 individuals (since this is the expected equilibrium density in the absence of parasites in the Beverton-Holt model) in each patch, with an initial parasite prevalence of 0.5. At initialisation, parasite virulence and host dispersal values were drawn from a uniform distribution taking values between 0 and 1, thus assuming standing genetic variation in the host and parasite populations. Table 1 summarises all model parameters and their descriptions.

**Table 1:** Model parameters (parameter values in the focal scenario are highlighted)

| Model Parameter/Variable | Description | Values |
| --- | --- | --- |
| $\mu$ | Dispersal mortality | **0, 0.01, 0.05, 0.1, 0.2, ..., 1** |
| $\lambda_0$ | Intrinsic growth rate of the host according to the Beverton-Holt model | **4** |
| $\alpha$ | Intra-specific competition coefficient of the host according to the Beverton-Holt model | **0.01** |
| $v$ | Parasite virulence acting on the fecundity of the host | **evolves;** 0.80, 0.82, ..., 1 |
| $d$ | Host dispersal probability | **evolves;** 0.01, 0.05, 0.1, 0.2, ..., 1 |
| $a$ | Searching efficiency of the parasite | **0.01** |
| $\beta_{max}$ | Maximum possible parasite transmission | **8** |
| $s$ | Shape of virulence-transmission relationship | **0.5** |
| $m_v$ | Mutation rate of parasite virulence | **0.01** |
| $m_d$ | Mutation rate of host dispersal probability | **0.01** |
| $\sigma_v$ | Mutation effect of parasite virulence (logit transform) | **0.5** |
| $\sigma_d$ | Mutation effect of host dispersal probability (logit transform) | **0.5** |

The main results focus on the equilibrium values corresponding to the co-evolution of host dispersal (evolving dispersal; ED) and parasite virulence (evolving virulence; EV) for different conditions of dispersal mortality *µ* = 0, 0.01, 0.05, 0.1, 0.2, …, 1. In order to better understand these results, we ran additional simulations, corresponding to the same conditions of dispersal mortality but fixing dispersal (fixed dispersal; FD) and allowing only virulence to evolve (EV) and vice versa (ED, FV).

## Results and discussion

### Effect of landscape structure on co-evolution of host dispersal and parasite virulence

Overall, when dispersal mortality is low, our model predicts that evolutionarily stable (ES) parasite virulence is greater (Fig. 1 A, C) and ES host dispersal is lower (Fig. 1 B, C) in riverine aquatic landscapes relative to terrestrial landscapes. However, when dispersal mortality is high, low dispersal and virulence evolve in both landscape types (Fig. 1 C). In Fig. 1 C, focusing on the relationship between ES virulence and ES dispersal, in co-evolutionary simulations, the following patterns are clear: There is an overall positive relationship between ES parasite virulence and ES host dispersal for both landscape types. In riverine aquatic landscapes, ES virulence increases with ES host dispersal reaching its maximal possible value of *v* = 1 at a median ES dispersal rate of *d* = 0.42 when dispersal mortality *µ* = 0. In contrast, a wider range of dispersal rates evolve in terrestrial landscapes (up to *d* = 0.97 when dispersal mortality *µ* = 0); however, virulence does not attain its maximal possible value but rather saturates beyond an ES host dispersal rate of *d* = 0.09, which is obtained at a dispersal mortality of *µ* = 0.8.

**Figure 1:**
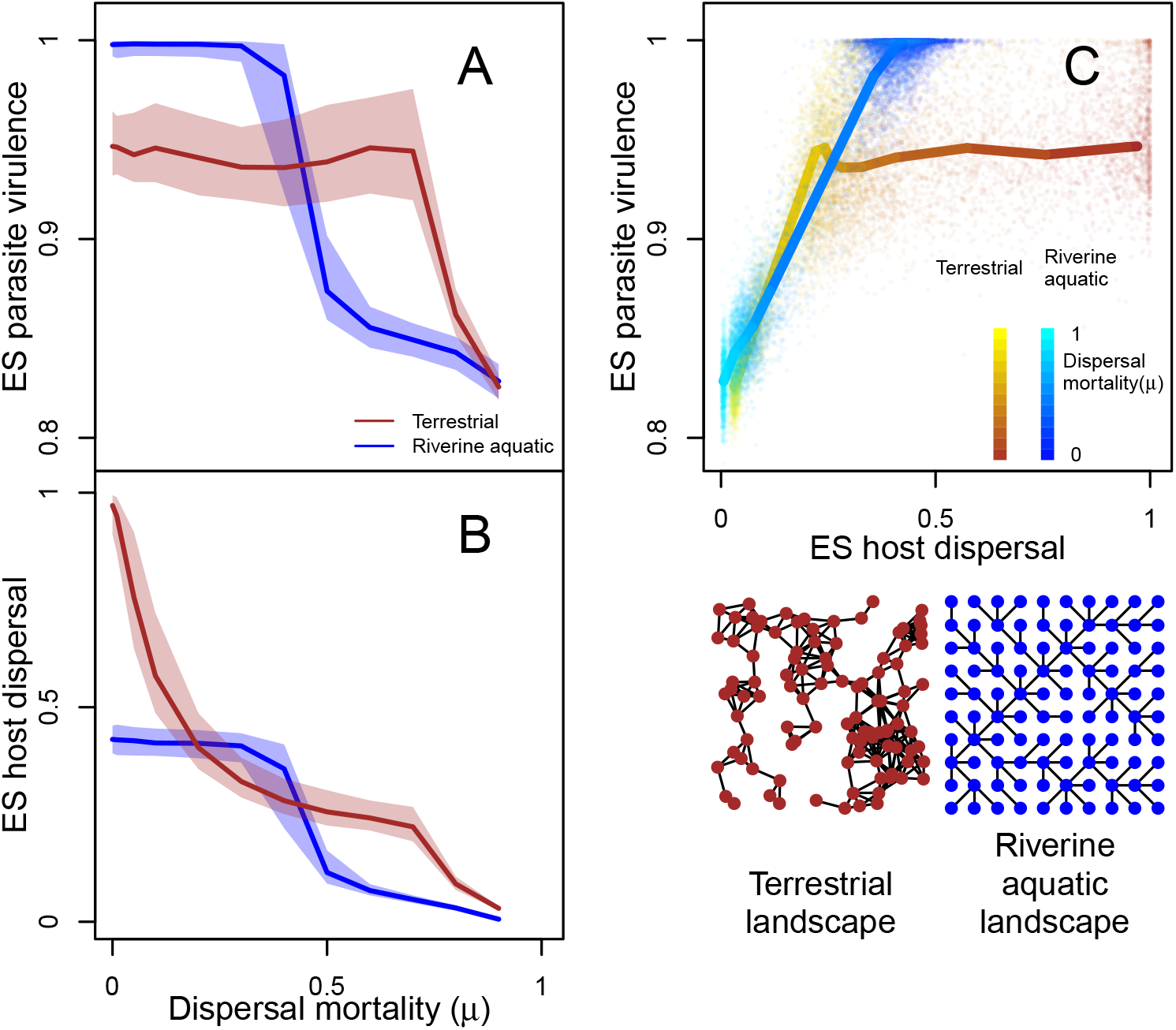
Evolutionarily stable (ES) parasite virulence and ES host dispersal as a function of dispersal mortality *µ* in riverine aquatic (modelled as optimal channel networks; OCNs) and terrestrial landscapes (modelled as random-geometric graphs; RGGs) when both dispersal and virulence can evolve (ED + EV). For each parameter combination, ES virulence and dispersal are represented by the median genotypic value of the trait over all individuals in the last time step of a simulation *t* = 5000. All measures are medians over 1000 landscape realisations. Solid lines in A and B are medians, and the shaded areas are interquartile ranges. A: ES parasite virulence as a function of dispersal mortality (*µ*). ES virulence decreases with dispersal mortality for both landscape types. Virulence is greater in riverine aquatic landscapes at low dispersal mortality and lower at high dispersal mortality relative to terrestrial landscapes. B: ES host dispersal rates as a function of dispersal mortality. In general, ES dispersal decreases with dispersal mortality, but when dispersal mortality is low, terrestrial landscapes lead to the evolution of greater dispersal relative to riverine aquatic landscapes. C: ES parasite virulence vs. ES host dispersal pooled across all values of dispersal mortality. The points indicate the evolved parasite virulence vs. evolved host dispersal for one landscape realisation, with the colour indicating the value of dispersal mortality. In terrestrial landscapes, a wider range of dispersal evolves, and virulence increases with dispersal. In riverine aquatic landscapes, virulence again increases with dispersal, but dispersal rates are relatively low. Fixed model parameters: intrinsic growth rate *λ*_0_ = 4, intra-specific competition coefficient *α* = 0.01, maximum possible transmission *β*_*max*_ = 8, shape of virulence-transmission function *s* = 0.5.

These results are markedly different from a previous similar model where host dispersal was not allowed to evolve, particularly for riverine aquatic landscapes (Deshpande et al., 2023). While a similar saturating relationship between ES parasite virulence and fixed host dispersal is obtained for terrestrial landscapes (see Fig. 2 F), in riverine aquatic landscapes a unimodal relationship is expected (see Fig. 2 A), which is absent when the two traits co-evolve (Fig. 1 C). Thus, since ES host dispersal is constrained to be under *d* = 0.42 in riverine aquatic landscapes, the combination of lower parasite virulence and high host dispersal cannot occur when the two traits co-evolve. This interplay between dispersal evolution and virulence evolution is further explored in the next section.

**Figure 2:**
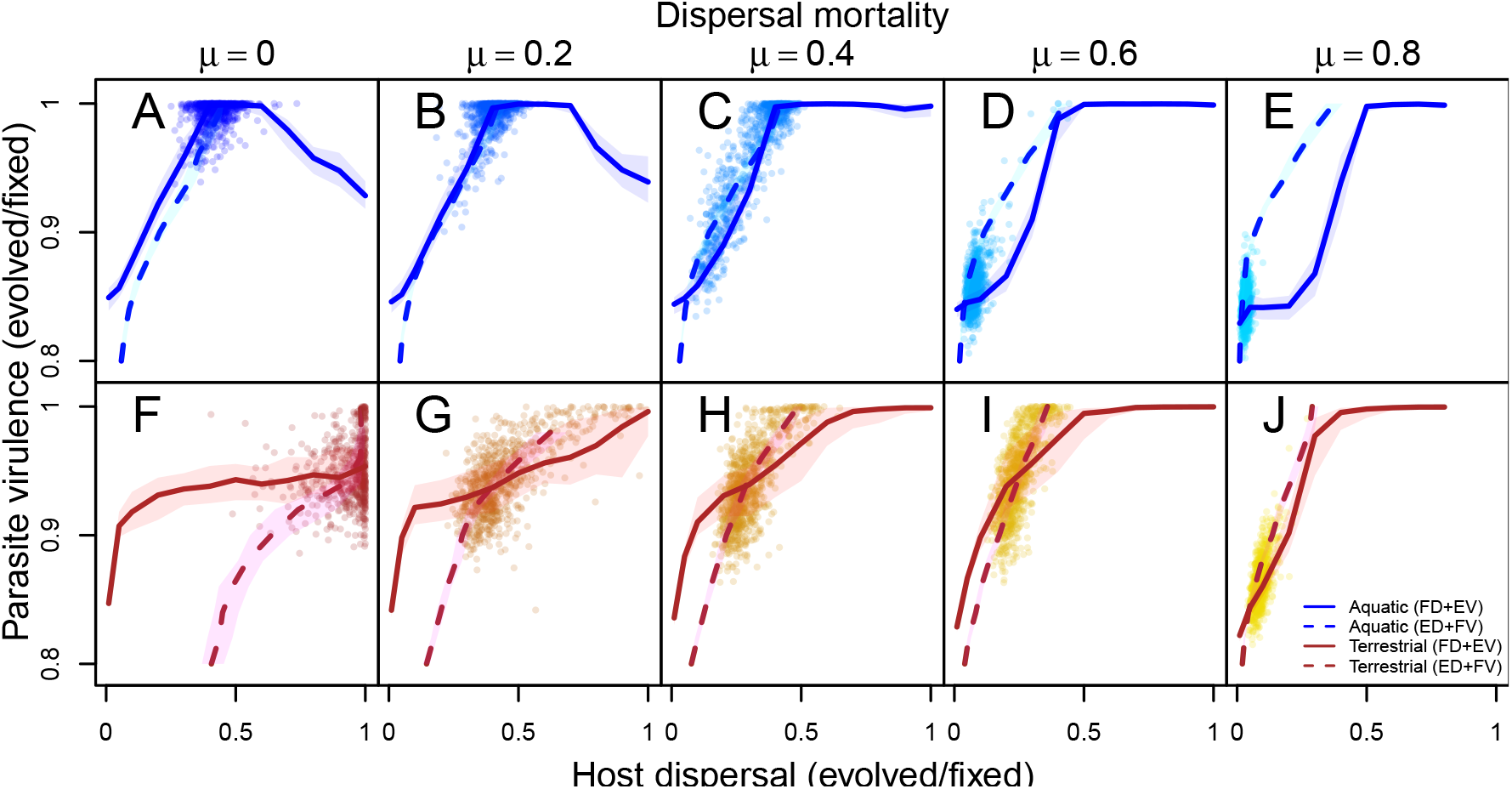
ES parasite virulence as a function of fixed host dispersal (FD + EV; solid lines), ES host dispersal as a function of parasite virulence (ED + FV; dashed lines), and ES parasite virulence as a function of ES host dispersal (ED + EV; points). From left to right, dispersal mortality increases, and the top rows show riverine aquatic landscapes, and the bottom rows the terrestrial landscapes. Fixed model parameters: intrinsic growth rate *λ*_0_ = 4, intra-specific competition coefficient *α* = 0.01, maximum possible transmission *β*_*max*_ = 8, shape of virulence-transmission function *s* = 0.5.

### Dispersal mortality modulates the effect of landscape structure on (co)-evolution of host dispersal and parasite virulence

Since dispersal mortality is clearly a modulator of the differences in evolved host dispersal and parasite virulence found between the two landscape types, we run additional simulations for varying dispersal mortality in which: 1) host dispersal is fixed and parasite virulence evolves; and 2) host dispersal evolves and parasite virulence is fixed. We compare these results to our focal scenario in which both traits evolve.

#### Low dispersal costs

When dispersal costs are low and when we allow only virulence to evolve, fixing dispersal (FD + EV; e.g., *µ* = 0 and *µ* = 0.2; Fig. 2 A–B, F–G solid lines), virulence is unimodal in aquatic landscapes and attains a maximum of *v* = 1 at intermediate dispersal, whereas in terrestrial landscapes, virulence saturates with increasing dispersal to a virulence *v <* 1. Deshpande et al. (2023) show that these differences in virulence evolution arise from the two landscapes generating different patterns of parasite relatedness, hence kin selection found in both landscapes. This is also consistent with previous work (O’Keefe and Antonovics, 2002; Wild et al., 2009; Lion and Boots, 2010) that highlights kin selection as a key mechanism in virulence evolution. Particularly, Deshpande et al. (2023) show that at intermediate dispersal rates, due to the heterogeneous degree distribution of riverine aquatic landscapes, the high-connectivity patches have large enough infected densities such that they never go extinct and, therefore, accumulate a larger neutral diversity of parasite strains, leading to low relatedness, which drives up ES virulence. This does not happen in terrestrial landscapes because of less heterogeneous degree distributions (Deshpande et al., 2023).

Further, in both landscapes, when virulence is fixed, dispersal evolves to increasing values with increasing virulence (ED + FV; e.g., *µ* = 0 and *µ* = 0.2; Fig. 2 A–B, F–G dashed lines), which is consistent with Chaianunporn and Hovestadt (2012b) who show that parasite-induced oscillations lead to local extinction of the host, which leads to selection for greater dispersal as a bet hedging strategy. We confirm this by showing that host extinctions increase as a function of virulence, leading to a corresponding increase in dispersal (Fig. S4). For a given fixed virulence, host dispersal always evolves to larger values in terrestrial as opposed to riverine aquatic landscapes. This is consistent with Henriques-Silva et al. (2015) and Fronhofer and Altermatt (2017), who show for a single species that riverine aquatic landscapes lead to the evolution of lower dispersal rates due to the fact that these landscapes create spatial heterogeneity in equilibrium population sizes. Generally, patches with low connectivity receive fewer dispersers than they send; thus, competition in these patches is low relative to high connectivity patches that receive more individuals than they send out. Since there are more degree-1 patches in riverine networks, this implies that hosts disperse to relatively more unfavourable (high competition conditions), which selects against dispersal. We show in Fig. S5 that this mechanism holds when there is the co-evolution of parasite virulence. However, due to the parasite impact on demography, reverse patterns of host density in which host density is greater in low connectivity patches are also possible at very low dispersal rates. However, since overall there is selection for dispersal to values greater than these extremely low dispersal rates, the patterns of host population density resemble those in Henriques-Silva et al. (2015) and Fronhofer and Altermatt (2017). Clearly, while the presence of a virulent parasite leads to overall increased dispersal rates in both landscape types, the effect of landscape structure leads to differential dispersal evolution for a given virulence.

Taken together, the heterogeneous structure of riverine aquatic networks leads to spatial heterogeneity in host-parasite densities, which leads to the evolution of lower dispersal rates relative to terrestrial landscapes. Further, this spatial heterogeneity also leads to lower parasite relatedness relative to terrestrial landscapes, leading to the evolution of greater parasite virulence relative to terrestrial landscapes.

#### High dispersal costs

When dispersal costs are high (e.g., *µ* = 0.4, 0.6, 0.8; Fig. 2 C–E, H–J), when we fix dispersal and allow virulence to evolve, we obtain low virulence for low dispersal and high virulence for high dispersal in both landscape types. Further, since dispersal is costly, it evolves to lower values in both landscape types when we fix virulence. Therefore, when dispersal is costly and both dispersal and virulence can evolve, dispersal is lower and virulence evolves to lower values in both landscape types.

## General discussion

Our study shows that landscape structure impacts the co-evolution of host dispersal and parasite virulence. Particularly, terrestrial landscapes are predicted to have more dispersive hosts and less virulent parasites compared to riverine aquatic landscapes, as long as dispersal mortality is low. When dispersal mortality is high, both riverine aquatic landscapes and terrestrial landscapes should have less dispersive hosts and less virulent parasites. More generally, the spatial heterogeneity in host-parasite densities generated by riverine aquatic landscapes (Deshpande et al., 2023) leads to both the evolution of greater virulence by allowing for lower parasite relatedness and the evolution of lower dispersal rates (Fronhofer and Altermatt, 2017), relative to terrestrial landscapes, which are less heterogeneous in their degree distribution.

### Implications of considering co-evolution of dispersal for trait evolution in complex landscapes

Previous work addressing the evolution of parasite virulence for simplified spatial structures identified kin selection as the key evolutionary driver (Wild et al., 2009; Lion and Boots, 2010). Extending this work to complex spatial structures, such as terrestrial and riverine aquatic landscapes, Deshpande et al. (2023) demonstrated the possible effects of network topology on parasite relatedness and virulence evolution. In the present study, we take this approach one step further by allowing concomitant dispersal and virulence evolution. One main finding is that this trait co-evolution constrains the range of possible evolutionary outcomes in the two landscapes: In terrestrial networks, the entire range of dispersal rates can evolve, but in riverine aquatic landscapes, only low dispersal rates are evolutionarily possible. Thus, while considering virulence evolution alone would lead us to predict that highly dispersive hosts and less virulent parasites are possible in riverine aquatic landscapes, allowing for co-evolution of host dispersal, we show that this is not possible.

In a more ecological context, it has been shown that the evolution of lower dispersal in riverine landscapes reinforces classical metapopulation dynamics of single species (Fronhofer and Altermatt, 2017). Our work reveals similar constraints imposed by spatial network structure on dispersal evolution in the presence of an antagonistic species. Thus, the same network features may not only be relevant for predicting ecological dynamics in multi-species communities, but also have potential knock-on effects on the evolution of traits mediating species interaction, such as virulence.

In conclusion, when studying the impact of spatial network structure on trait evolution, dispersal evolution should be taken into account. For example, in a host-parasite context, dispersal evolution may be relevant for studying the impact of landscape structure (Jousimo et al., 2014) on host-parasite co-evolution and local adaptation (Gandon et al., 1996). We speculate that the evolution of lower dispersal rates in riverine aquatic landscapes might lead to local adaptation for a larger parameter space, for example.

### Dispersal evolution in a multi-species context: role of biotic factors and landscape structure

As our results show, when trait evolution feeds back on the evolution of dispersal, considering the evolution of both traits is critical (Zilio et al., 2024). The ultimate causes of dispersal evolution in a single species context are well understood from a theoretical perspective (Bowler and Benton, 2005; Duputié and Massol, 2013). In a multi-species context, these mechanisms likely remain the same (Chaianunporn and Hovestadt, 2012b; Fronhofer et al., 2024) and traits mediating interspecific interaction (such as virulence) impact the evolution of dispersal via its effect on demography and genetic structure (its impact on kin competition or inbreeding depression). These biotic interactions generate conditions that can favour the evolution of dispersal (e.g., by inducing extinction-recolonisation dynamics Chaianunporn and Hovestadt 2012b) or impose additional costs by introducing spatial heterogeneity (Drown et al., 2013). The exact impact of species interaction strength on dispersal evolution may depend on the interaction type (Fronhofer et al., 2024) and its exact impact on demography and genetic structure. Thus, Zilio et al. (2024) argue in the case of antagonistic interactions, such as host-parasitoid, host-parasite, and predator-prey systems, that when demographic impacts are large, co- and concurrent evolution of interaction traits and dispersal will be relevant.

The precise effect of these interactions has also been shown to depend on the abiotic context. For example, by modelling the joint evolution of niche width and dispersal in host-parasite systems, Chaianunporn and Hovestadt (2012a) have shown that heterogeneity in external abiotic conditions selects against both host and parasite dispersal. Our work contributes two important aspects: First, we show that while increased parasite virulence generally leads to dispersal rates that increase with virulence, the exact relationship depends on the landscape context. This shows that even without an external environmental factor (e.g., temperature), landscape structure alone can generate diverging conditions between ecological systems. Second, while previous studies fixed the values of virulence and searching efficiency (Chaianunporn and Hovestadt, 2012b), the dispersal rates evolved here are constrained by the possible values of virulence, which are also driven by landscape structure.

Finally, it should be noted that our study assumes that parasites only travel with their dispersing hosts. Although this is common in many biological systems, it is also possible that parasite dispersal evolves as a trait in its own right. This latter case has been studied in certain models of host-parasite (Chaianunporn and Hovestadt, 2012b) and host-parasitoid systems (Green, 2009), and they show that parasite dispersal rates also tend to increase with virulence. Thus, since the demographic impacts are tightly coupled, we speculate that our results would remain unchanged if we allowed for parasite dispersal to evolve independently. However, future research may study the impact of different dispersal networks for hosts and parasites and possible trade-offs between virulence and dispersal (Osnas et al., 2015; Nørgaard et al., 2021).

### Landscape structure as a driver of eco-(co)-evolutionary dynamics

The importance of spatial network structure in driving ecological dynamics is increasingly being recognised (Savary et al., 2023). Metapopulation (Fronhofer and Altermatt, 2017) and metacommunity (Carrara et al., 2012) dynamics, range expansions (Rayfield et al., 2023) and the spread of perturbations (Gilarranz et al., 2017) have all been shown to depend on spatial network structure. As we show here, spatial network structure also has the potential to drive co-evolutionary interactions in species communities. We illustrate this for the co-evolution of parasite virulence and host dispersal, showing: 1) direct effects of landscape structure on virulence evolution; 2) indirect effects of landscape structure on virulence via its effect on dispersal evolution; and 3) direct effects of dispersal rates on virulence along with how the trait feeds back on dispersal evolution. In our model, the key network feature is the variation in patch connectivity (degree distribution): Particularly, the highly heterogeneous degree distribution of riverine landscapes relative to terrestrial landscapes leads to spatial heterogeneity in host and parasite densities. This variation produces both the observed patterns of parasite relatedness leading to evolution of greater virulence, as shown in Deshpande et al. (2023), and leads to evolution of lower dispersal rates, as shown here and previously (Fronhofer and Altermatt, 2017). Our work underlines the importance of dispersal as a key eco-evolutionary linker trait (Govaert et al., 2019; Fronhofer et al., 2023) that has to be considered within its respective spatial network.

## Author contributions

JND and EAF conceived the study. JND and RSP developed and analysed the models in collaboration with EAF. JND wrote the manuscript in collaboration with EAF and all authors commented on the draft.

## Acknowledgements

This work was funded by a grant from the Agence Nationale de Recherche to O.K. (grant no. ANR-20-CE02-0023-01). This is publication ISEM-YYYY-XXX of the Institut des Sciences de l’Evolution – Montpellier.

## Data availability

Model code is available via GitHub and a Zenodo DOI: https://doi.org/10.5281/zenodo.13992404.

## Notes

https://doi.org/10.5281/zenodo.13992404

### Competing Interest Statement

The authors have declared no competing interest.

