## Supplementary information for "Co-evolution of host dispersal and parasite virulence in complex landscapes"

### Supplementary figures

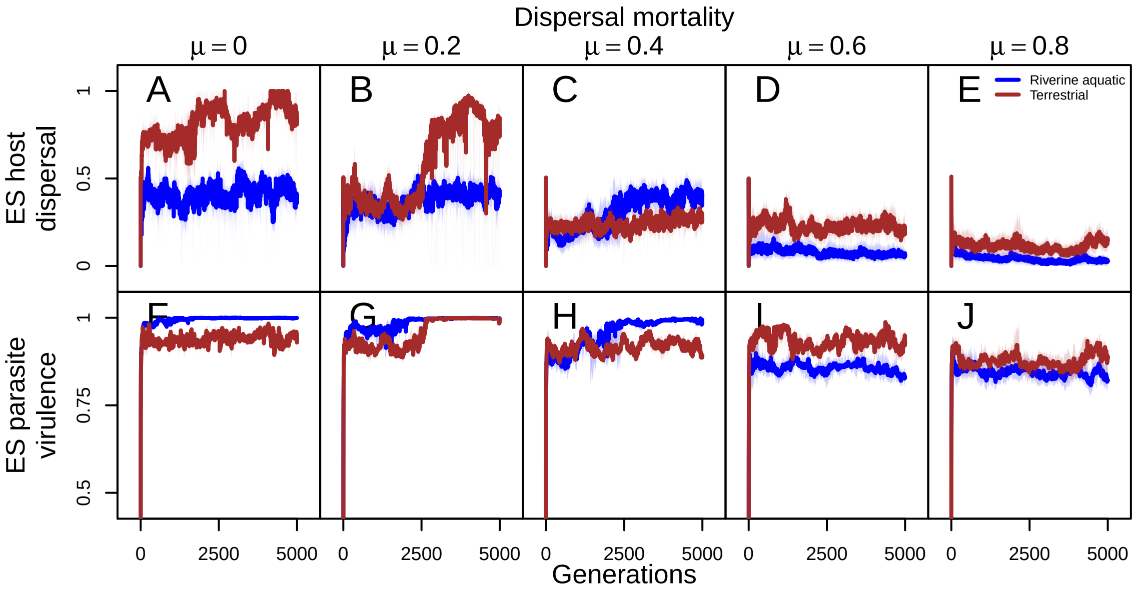


Figure S1: Co-evolutionary dynamics of host dispersal and parasite virulence in terrestrial and riverine aquatic landscapes. From left to right, dispersal mortality increases. A–E: Host dispersal as a function of time averaged over all patches for one representative replicate simulation. F–J: Parasite virulence as a function of time averaged over all patches. Fixed model parameters: intrinsic growth rate $\lambda_0 = 4$, intra-specific competition coefficient $\alpha = 0.01$, maximum possible transmission $\beta_{max} = 8$, shape of virulence-transmission function $s = 0.5$.

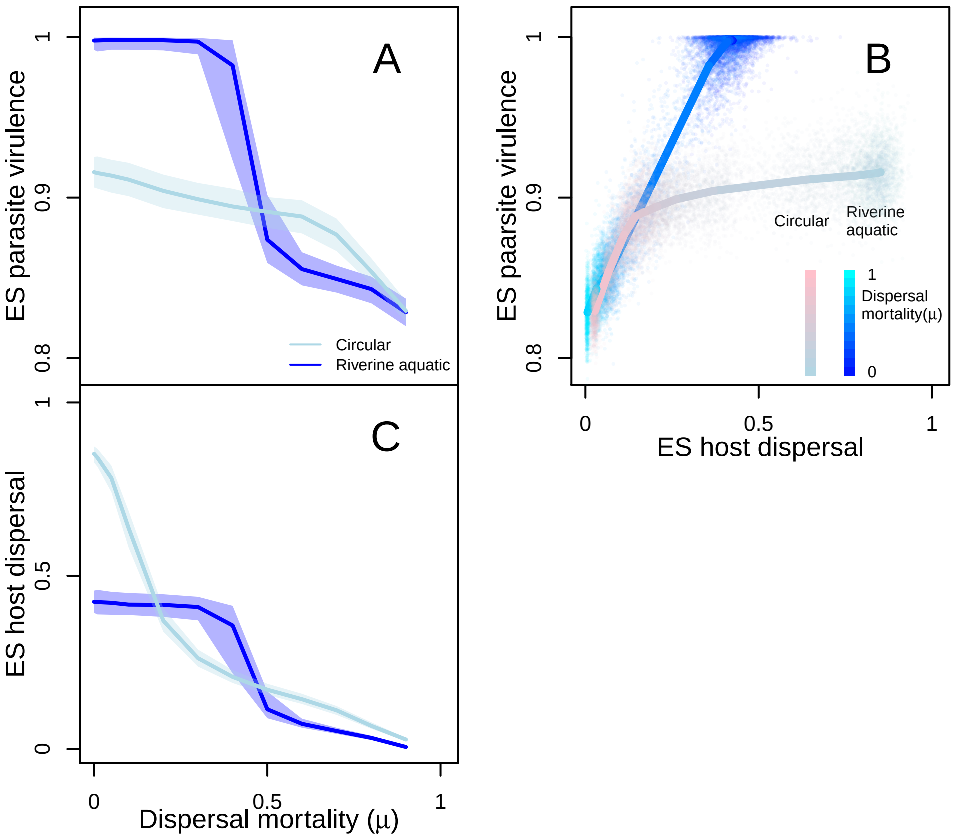


Figure S2: Comparing the co-evolution of host dispersal and parasite virulence in aquatic landscapes (modelled by optimal-channel networks of average degree of 1.98), to circular networks, which are regular networks with each patch having 2 neighbours. Evolutionarily stable (ES) parasite virulence and ES host dispersal as a function of dispersal mortality $\mu$ in aquatic landscapes and circular networks when both dispersal and virulence can evolve (ED + EV). For each parameter combination, ES virulence and dispersal are represented by the median genotypic value of the trait over all individuals in the last time step of a simulation $t = 5000$. All measures are medians over 1000 landscape realisations for riverine aquatic landscapes and 1000 replicate simulations for circles. Solid lines in A and B are medians and the shaded areas are interquartile ranges. A: ES parasite virulence as a function of dispersal mortality ($\mu$). ES virulence decreases with dispersal mortality in both landscape types, but riverine aquatic landscapes attain a higher ES virulence compared to circular networks. B: ES host dispersal rates as a function of dispersal mortality. ES dispersal decreases with dispersal mortality in both landscape types, with circular landscapes attaining higher host dispersal rates when dispersal mortality is low. C: ES parasite virulence vs. ES host dispersal pooled across all values of dispersal mortality. The points indicate the evolved parasite virulence vs. evolved host dispersal for one landscape realisation, with the colour indicating the value of dispersal mortality. Overall ES virulence increases with ES dispersal in both landscape types, however, circular networks lead to lower virulence and greater dispersal overall relative to riverine aquatic landscapes. This indicates that the characteristic pattern of co-evolution of host dispersal and parasite virulence in riverine aquatic landscape depends on higher order network properties, rather than average degree alone. Fixed model parameters: intrinsic growth rate $\lambda_0 = 4$, intra-specific competition coefficient $\alpha = 0.01$, maximum possible transmission $\beta_{max} = 8$, shape of virulence-transmission function $s = 0.5$.

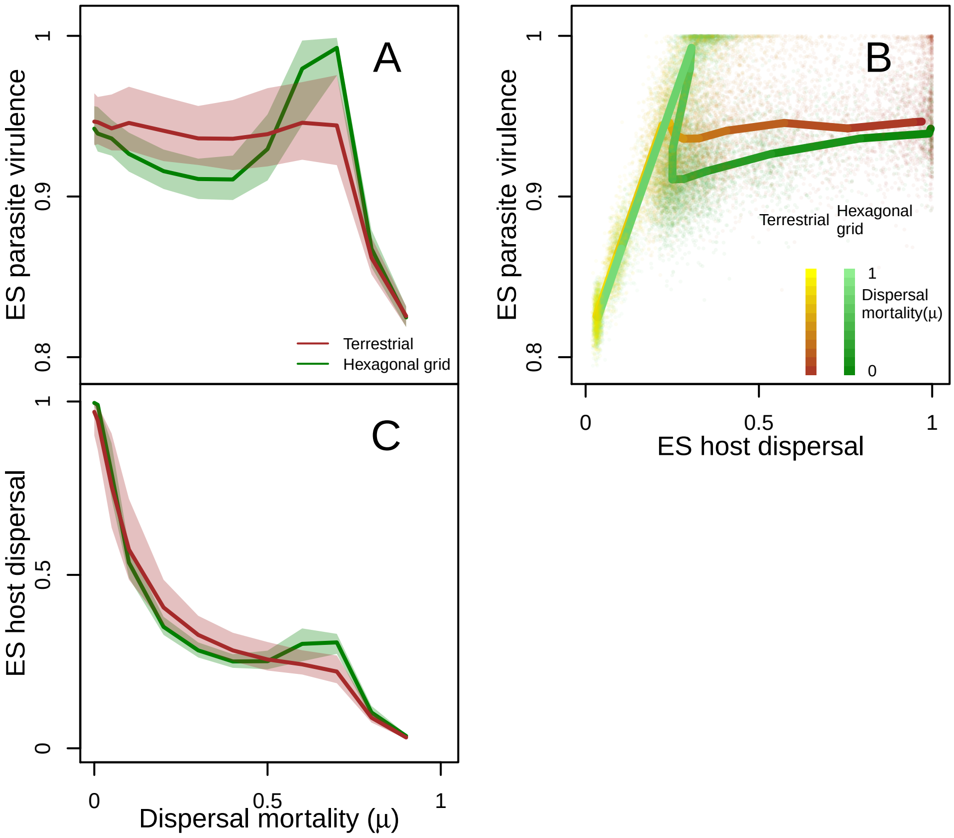


Figure S3: Comparing the co-evolution of host dispersal and parasite virulence in terrestrial landscapes (modelled by random-geometric graphs of average degree of 6), to hexagonal grids, which are regular networks with each patch having 6 neighbours. Evolutionarily stable (ES) parasite virulence and ES host dispersal as a function of dispersal mortality $\mu$ in terrestrial landscapes and hexagonal grids when both dispersal and virulence can evolve (ED + EV). For each parameter combination, ES virulence and dispersal are represented by the median genotypic value of the trait over all individuals in the last time step of a simulation $t = 5000$. All measures are medians over 1000 landscape realisations for terrestrial landscapes and 1000 replicate simulations for hexagonal grids. Solid lines in A and B are medians and the shaded areas are interquartile ranges. A: ES parasite virulence as a function of dispersal mortality ($\mu$). ES virulence decreases with dispersal mortality for terrestrial landscapes, but peaks at higher dispersal mortality in hexagonal grids. B: ES host dispersal rates as a function of dispersal mortality. In terrestrial landscapes, ES dispersal decreases with increasing dispersal mortality but in hexagonal grids, there is a an initial decrease in ES dispersal, followed by a small increase and then decreases. C: ES parasite virulence vs. ES host dispersal pooled across all values of dispersal mortality. The points indicate the evolved parasite virulence vs. evolved host dispersal for one landscape realisation, with the colour indicating the value of dispersal mortality. In terrestrial landscapes and hexagonal grids, the entire range of dispersal rates evolve, however, virulence reaches higher values in hexagonal grids, before decreasing again. This result is consistent with Deshpande et al. (2023) and shows that even though the two landscapes have similar average degrees, patterns of co-evolution of parasite virulence and host dispersal differ between them, highlighting the importance of considering higher order network properties. Fixed model parameters: intrinsic growth rate $\lambda_0 = 4$, intra-specific competition coefficient $\alpha = 0.01$, maximum possible transmission $\beta_{max} = 8$, shape of virulence-transmission function $s = 0.5$.

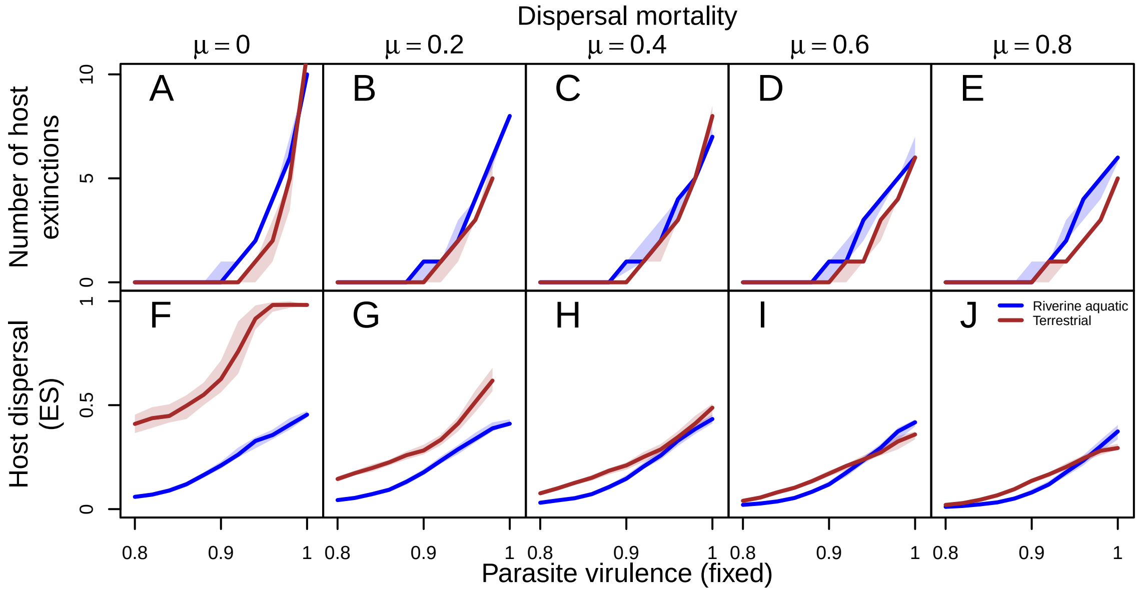


Figure S4: Number of host extinctions and evolved host dispersal in simulations with fixed virulence and dispersal evolving (ED+FV). From left to right, dispersal mortality increases. A–E: Number of host extinctions as a function of virulence in simualtions where dispersal can evolve but virulence is fixed. At each time step, we track the number of patches that were previously occupied that go extinct, this is termed as number of host extinctions. Clearly, the number of host extinctions increases with parasite virulence consistent with Chaianunporn and Hovestadt (2012) due to higher amplitude oscillations in host-parasite densities. F–J: Host dispersal probability as a function of virlulence in ED+FV simulations. Again, consistent with Chaianunporn and Hovestadt (2012), ES dispersal increases with virulence. Fixed model parameters: intrinsic growth rate $\lambda_0 = 4$, intra-specific competition coefficient $\alpha = 0.01$, maximum possible transmission $\beta_{max} = 8$, shape of virulence-transmission function $s = 0.5$.

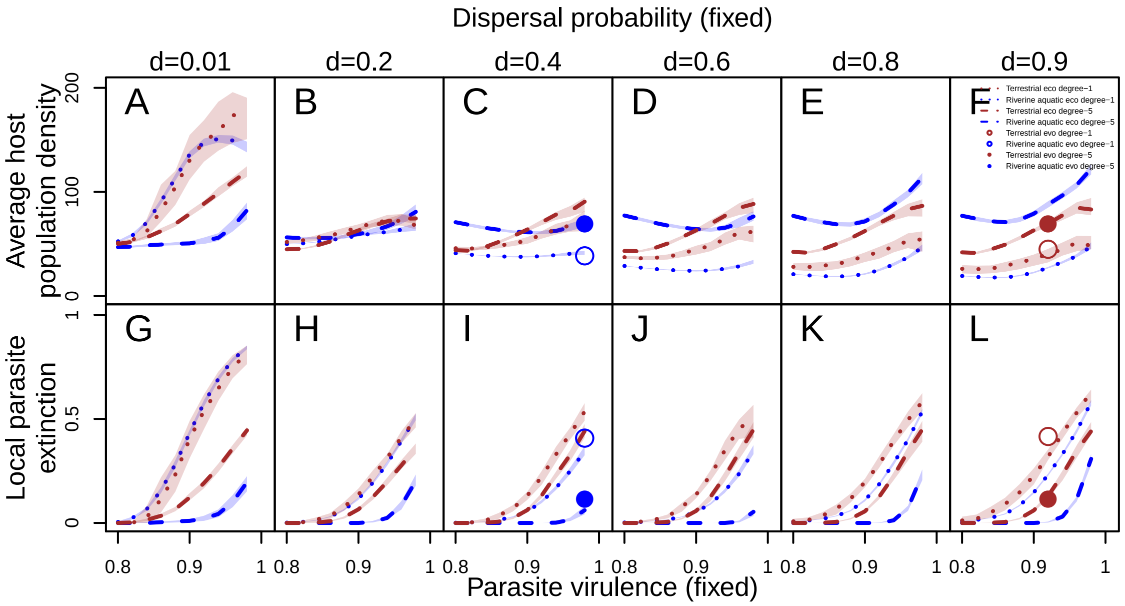


Figure S5: Spatial heterogeneity in host population density and local parasite extinction as a function of fixed virulence, for varying fixed dispersal rates (lines) and dispersal mortality $\mu = 0$ and for simulations in which both dispersal and virulence can evolve (points) for terrestrial and riverine aquatic landscapes. From left to right dispersal probability increases. A–F: Average host population density in degree-1 (dotted line) and degree-5 (dashed line) patches as a function of ecologically fixed virulence and dispersal. The points indicate average host population density in simulations where both dispersal and virulence can evolve. G–L: Local parasite extinction or the fraction of patches in which the parasite is absent in degree-1 and degree-5 patches as a function of virulence. The points indicate local parasite extinction in simulations where both dispersal and virulence can evolve. Clearly, given a dispersal rate the difference in host population density between degree-1 and degree-5 patches is greater in riverine aquatic landscapes relative to terrestrial landscapes. However, at low dispersal rates host density is greater in degree-1 patches but as dispersal increases, this is reversed with greater host density in degree-5 patches for both landscape types. This is because there are two mechanisms at play: 1) following extinction of parasite in a patch, greater dispersal should lead to immediate recolonisation, thus at low dispersal rates the failure of the parasite to recolonise patches especially in low connectivity patches allows the host to grow to greater population density and 2) greater dispersal increases the asymmetry in population density between patches, with low connectivity patches sending out more disperses than they receive (Fronhofer and Altermatt, 2017). This leads to greater host population density in high connectivity patches. Further, low dispersal mortality $\mu = 0$ and parasitism (Chaianunporn and Hovestadt, 2012) create conditions in which high dispersal can evolve, thus at these dispersal rates, host population density is greater in high connectivity patches as seen in the evolutionary simulations. Thus, in riverine aquatic landscapes, if dispersal were to evolve to greater values than what we find, this would lead to greater net dispersal into high connectivity patches in which individuals face both higher intra-specific competition, ad a greater risk of being infected. Fixed model parameters: intrinsic growth rate $\lambda_0 = 4$, intra-specific competition coefficient $\alpha = 0.01$, maximum possible transmission $\beta_{max} = 8$, shape of virulence-transmission function $s = 0.5$, dispersal mortality $\mu = 0$.
